# Cloud-native distributed genomic pileup operations

**DOI:** 10.1101/2022.08.27.475646

**Authors:** Marek Wiewiórka, Agnieszka Szmurło, Paweł Stankiewicz, Tomasz Gambin

## Abstract

**Motivation:** Pileup analysis is a building block of many bioinformatics pipelines, including variant calling and genotyping. This step tends to become a bottleneck of the entire assay since the straightforward pileup implementations involve processing of all base calls from all alignments sequentially. On the other hand, a distributed version of the algorithm faces the intrinsic challenge of splitting reads-oriented file formats into self-contained partitions to avoid costly data exchange between computation nodes.

**Results:** Here, we present a scalable, distributed, and efficient implementation of a pileup algorithm that is suitable for deploying in cloud computing environments. In particular, we implemented: (i) our custom data-partitioning algorithm optimized to work with the alignment reads, (ii) a novel and unique approach to process alignment events from sequencing reads using the MD tags, (iii) the source code micro-optimizations for recurrent operations, and (iv) a modular structure of the algorithm. We have proven that our novel approach consistently and significantly outperforms other state-of-the-art distributed tools in terms of execution time (up to 6.5x faster) and memory usage (up to 2x less), resulting in a substantial cloud cost reduction. SeQuiLa is a cloud-native solution that can be easily deployed using any managed Kubernetes and Hadoop services available in public clouds, like Microsoft Azure Cloud, Google Cloud Platform, or Amazon Web Services. Together with the already implemented distributed range joins and coverage calculations, our package provides end-users with an unified SQL interface for convenient analyzing of population-scale genomic data in an interactive way.

**Availability:** https://biodatageeks.github.io/sequila/

**Supplementary information:** Supplementary data are available at *Bioinformatics* online.

## 1 Introduction

The sorted collection of the aligned sequencing reads can be transformed into a set of pileup records, also known as a coverage position summary. This format summarizes information about the base calls in all genomic positions from the reads aligned to a reference sequence, including total depth of coverage, non-reference (alternative) bases, and base qualities. Pileup format was designed to provide the evidence of the single-nucleotide variants or the short insertions/deletions at a given genomic position. It is commonly used as an entry point to the well-established variant calling pipelines (Li, 2011) as well as to novel approaches to variant detection frameworks based on the neural networks (Luo *et al*., 2020) or other methods e.g. the binomial model, partial-order alignment, and de Bruijn graph local assembly (Liu *et al*., 2021) in fast variant calling. Coverage position summary is also used for identification of somatic mutations and copy number variation (Koboldt *et al*., 2012).

Samtools suite (Li *et al*., 2009) includes the mpileup tool, a golden standard for both data format and correctness of pileup calculations; however it is a single-threaded program that does not provide the scalability feature (Sater *et al*., 2020).

Research development in the bioinformatics field emphasises a common need to use a technology that allows distributing long-lasting big data tasks into the multiple computing nodes or in the cloud computing infrastructure (Yuan and Wildish, 2020). In the recent Genome Analysis Toolkit (GATK, McKenna *et al*., 2010) version, several programs (including pileup calculations) have been implemented in a distributed manner ready to be run on the Apache Spark cluster. Other research studies confirm that big data programming paradigms can be successfully applied to many genomics analyses (Guo *et al*., 2018, Capuccini *et al*., 2020, Wiewiórka *et al*., 2018, Wiewiórka *et al*., 2017) including variant calling(Ahmad *et al*., 2021). The analysis of the ever-increasing genomic data sets involves significant financial investments and administrative efforts to maintain secure and fault-tolerant storage solutions as well as fast and scalable processing units. To minimize those efforts medical clinics and research centers consider migrating bioinformatics pipelines and custom analyses to private or public cloud infrastructure. The evolution towards cloud architecture is embraced by the widely used bioinformatics products both open-source (e.g. GATK) and commercial (e.g. DNA Nexus, Terra) (Koppad *et al*., 2021).

Although significant progress has been made, there are still areas in bioinformatics analyses that are not easily transferable to the distributed and cloud environments with a traditionally used sets of tools. In this article, we introduce an updated version of the SeQuiLa package that includes a cloud-native, scalable and extensible software for summarizing reads in a pileup format.

## 2 Materials and Methods

### 2.1 Rationale

The foundations for the distributed pileup algorithm are based on three key observations.

Firstly, the majority of bases in the aligned sequencing reads are concordant with the reference sequence. Therefore, we designed our algorithm to use both CIGAR strings and MD tags to handle deletions, insertions, and substitutions to avoid decoding and parsing the entire read sequence and base qualities. To the best of our knowledge, it is the only algorithm that takes advantage of this information for pileup construction.

Secondly, our pileup computation is divided into four units of work: (i) coverage computation, (ii) identification of non-reference base calls, (iii) collection of base qualities, and (iv) output projection. This decomposition allows us to reduce computational complexity by skipping certain steps that are not required.

Finally, the most limiting factor of performance and scalability for any distributed processing is the data exchange among the worker nodes that always requires costly data serialization as well as network transfer. Therefore, we proposed a new data partitions coalescing mechanism, which guarantees proper handling of reads overlapping more than one partition without the need of data shuffling. In addition, we use both BAM indexes for efficient partition boundaries adjustment and thus significantly reducing input/output (IO) operations.

### 2.2 Algorithm

#### 2.2.1 Defining partitions

Consider input sorted collection of the aligned sequencing reads *R* divided into *n* partitions by underlying file system (Fig 1A).

**Fig. 1.**
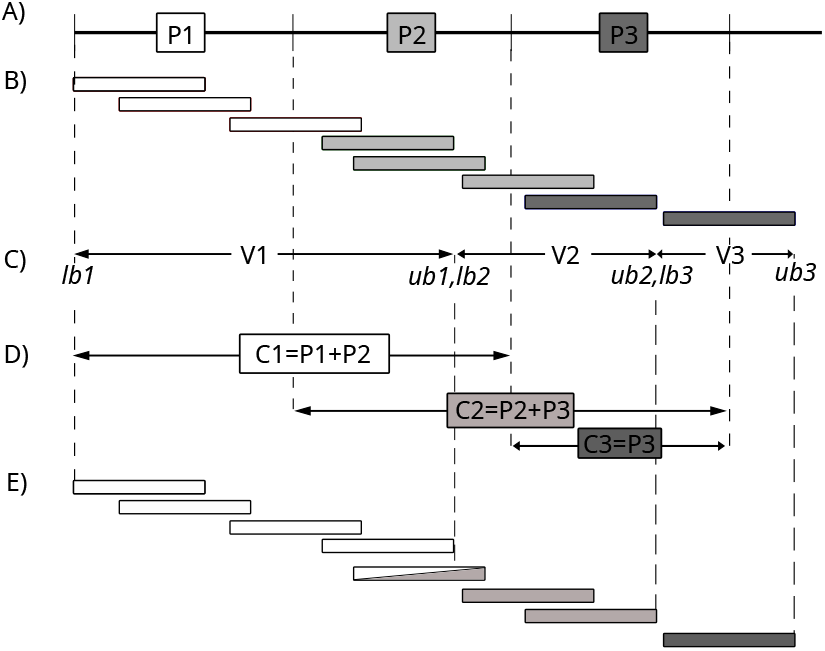
SeQuiLa partitioning algorithm: original distributed partitions (panel A); read assignment (color coded) to original partitions according to alignment starting position (panel B); virtual partitions and their boundaries calculated by algorithm 1 (panel C); coalesced partitions (panel D); read assignment (color coded) to coalesced partitions and corresponding virtual partitions (panel E). Note that some of the reads will be processed in more than one coalesced partition. This approach produces on average equally-sized virtual partitions (no data skewness) except for the first one and last one that are a bit larger and smaller than the rest, respectively.

For each partition we calculate two values: lower and upper bounds that create self-contained virtual read partitions (*V*1 = *L*1 − *U*1, *V*2 = *L*2 − *U*2, etc, from Fig 1C, algorithm 1). For clarity, in pseudo-code we assume that all reads are aligned to a single chromosome.

This information is further required for changing the default Apache Spark partitioning schema (*P*1, *P*2, …, *PN* - see Figure 1A), creating new coalesced partitions (*C*1, *C*2, …, *CN* - see Figure 1D). Within each coalesced partition the algorithm processes only the reads overlapping with the corresponding virtual partition (see Figure 1E). Note that our approach is very lightweight – it dynamically calculates boundaries for virtual partitions, which can be considered as view upon original Spark partitions. Unlike GATK which divides input data into fixed length multikilobase-size pieces called *shards*), we do not introduce any other level of data splitting and push all computations down to the partition level.

##### Algorithm 1 Calculating lower(*lb*) and upper(*ub*) bounds for self-contained read partitions

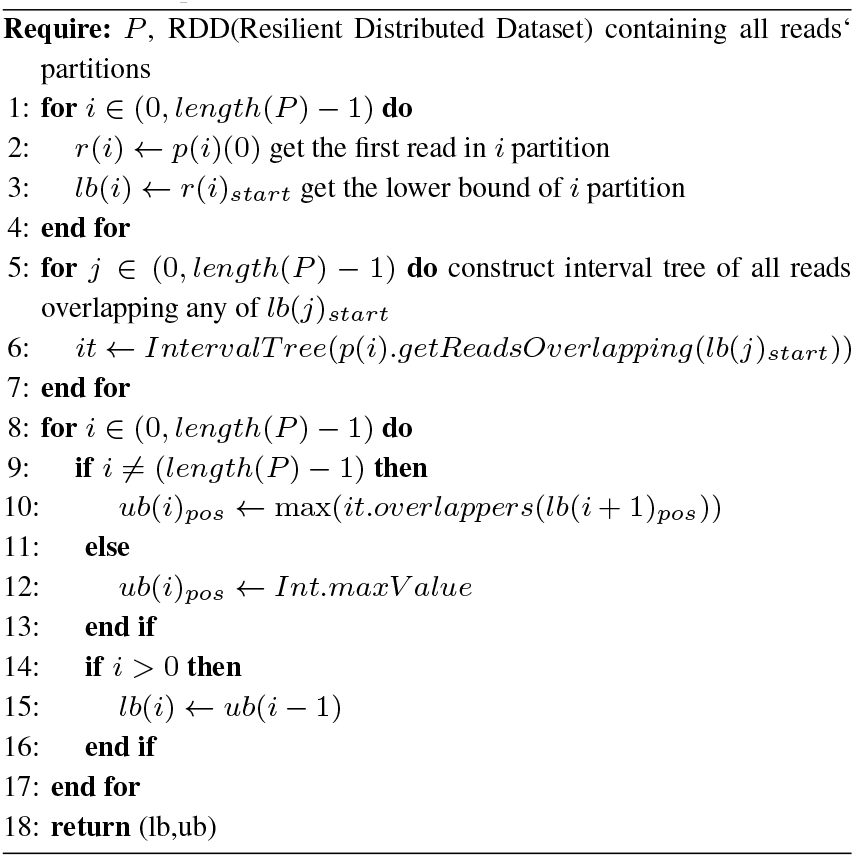

##### Algorithm 2 Calculating coverage and alternative alleles

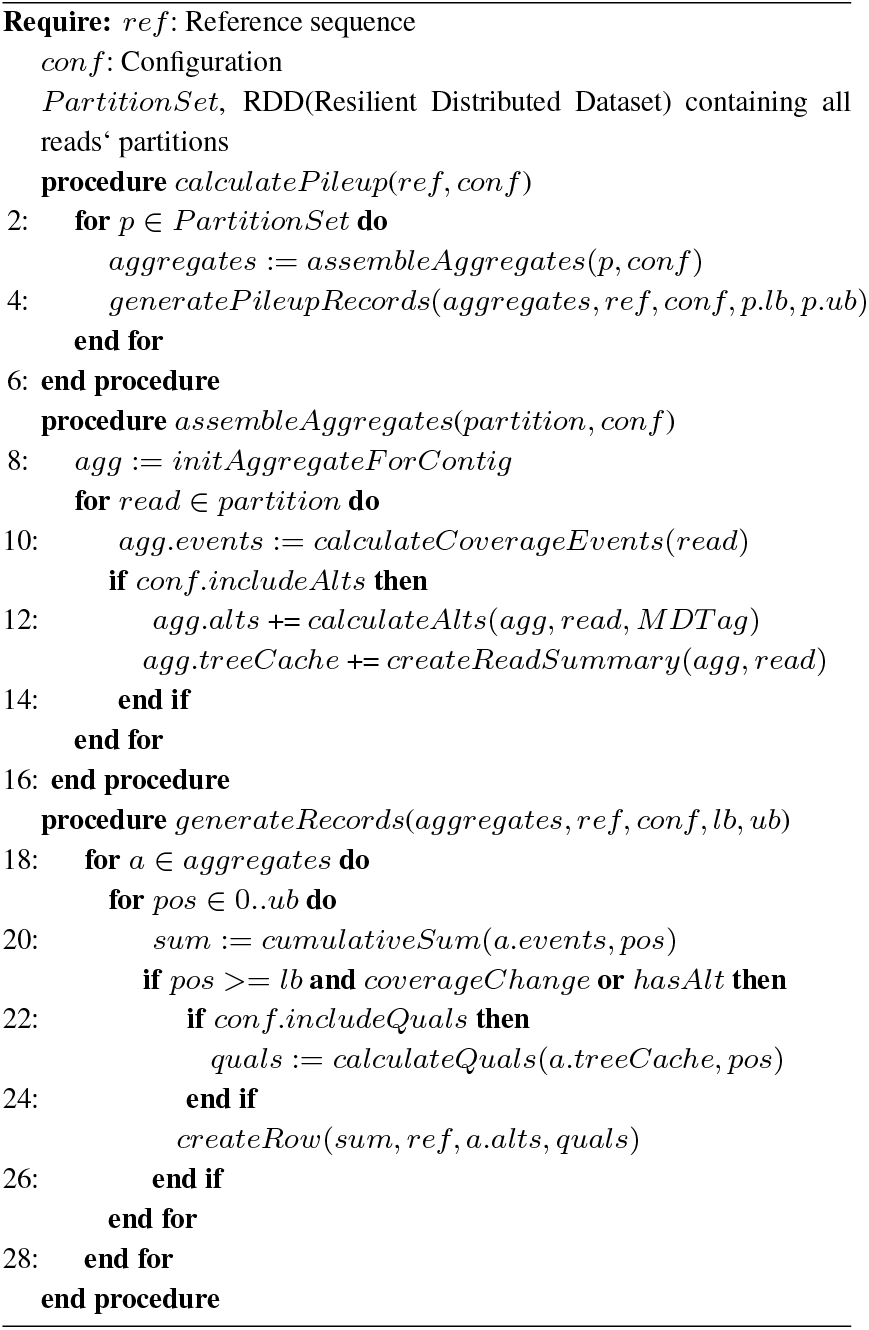

#### 2.2.2 Calculating coverage and alternative alleles

For each virtual partition the program generates an aggregate object holding: (i) an array of alignment events (i.e., start and end of alignment) which is gathered for the event-based coverage calculations, (ii) a map of alternative bases count calculated using read MD tag, and (iii) an interval tree structure of succinct read representation - *ReadSummary* (i.e., start, end, CIGAR derived configuration) used for further base qualities calculations. The program calculates base qualities only for the positions where at least one alternative base is present (2).

#### 2.2.3 Merging and rendering the results

Once all reads in each virtual partition are analyzed, the program calculates the set of final pileup records. Depending on configuration, it can generate different outputs: coverage only, coverage with alternative bases counts or the full pileup with coverage, alternative bases and base qualities.

### 2.3 Technical design

Our algorithm is implemented as a plugin to Apache Spark Catalyst optimizer (Armbrust *et al*., 2015). We used its three extension points: (i) SQL Analyzer - to register new table-valued functions, (ii) Planner - to add our optimized execution strategies for pileup calculations, and (iii) Logical Optimizer - to detect *CreateDataSourceTableAsSelectCommand* and *InsertIntoHadoopFsRelationCommand* actions and apply optimizations for direct vectorized writes into the Optimized Row Columnar (ORC) files (Figure 2).

**Fig. 2.**
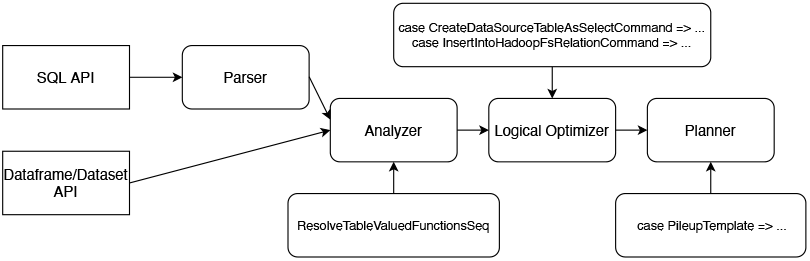
SeQuiLa extensions to Apache Spark Catalyst optimizer

We designed the relational model to represent alignments and pileup function results as proposed in Sun *et al*., 2018 and Smith *et al*., 2021. Our package provides both SQL (Structured Query Language) and Dataframe programming interfaces for the Scala and Python (https://github.com/biodatageeks/pysequila) languages.

For reading Binary Alignment Map (BAM) and CRAM files our solution can use Hadoop-BAM (Niemenmaa *et al*., 2012) or disq libraries as configured by the end-user. For better support of the CRAM files that have been recently added to the HTSJDK library, we extended the Hadoop-BAM project (https://github.com/biodatageeks/Hadoop-BAM). Also, minor changes required for serialization of genomic intervals parameters have been added to the disq (https://github.com/mwiewior/disq) library. For saving output we support not only ORC but also Parquet file format (Ivanov and Pergolesi, 2020). In our code, we re-used partition coalescing mechanism as implemented in the GATK.

### 2.4 Essential optimizations

Our main goal was to deliver the fast, distributed, and scalable implementation of a pileup algorithm. In additional to the already presented novel algorithm, we highlight other essential implementation decisions that improve overall software performance. They can be grouped into three main categories: (i) optimization of distributed processing, (ii) Scala source code micro-optimizations, and (iii) output vectorization and fine-tuning.

#### 2.4.1 Optimization of distributed processing

In the straightforward approach where the default partitioning is used, pileup implementation in Apache Spark would to be split into two stages with a data shuffle step in between. This would require either explicit caching of the intermediate results from the first one or at least partial recomputing of the evicted partitions to get the final results. Implementation of a custom partitioning mechanism (algorithm 1) to appropriately split data and determine the boundaries for each split was essential to achieve a single-pass solution without any extra data exchange between the executors or caching intermediary results.

#### 2.4.2 Source code micro-optimizations

We have observed an apparent speedup when using an interval tree to store a short representation of reads (*ReadSummary*) since data retrieval from this structure is performed frequently with interval conditions. After analyzing the profiling results in a form of flame graphs obtained with *async-profiler* (Nisbet *et al*., 2019), we identified the most time-consuming and frequently invoked methods, such as calculation of the relative position in a read for a given genomic coordinate, and re-implemented them in the state-aware manner, thus eliminating traversing collection on each call. Similarly, we substituted computationally expensive CIGAR parsing and interpretation with fast lookups to lazily evaluated custom objects with derived cigar configuration with quick checks for existence of clip (and its length) or deletion, as well as deletion and insertion positions.

#### 2.4.3 Output and auxiliary optimizations

The default output generation mechanism, which accounted for around 30% of the total processing time of our algorithm turned out to be another bottleneck. Therefore, we have implemented two novel approaches for optimizing output rendering.

First, we have developed a custom direct pileup record projection to Apache Spark’s internal binary row representation applying several micro optimizations e.g. casting reference bases and contig names to bytes and caching them in map to avoid repeating this task for each record. This mechanism can be used for further processing pileup rows within Spark-SQL engine as well as for persisting the results in any supported file format.

The second method is intended for saving the results in ORC file format only. Inspired by the idea of *direct-path load* introduced in relational database management systems, in particular in Oracle database (Heller, 2019), we implemented a mechanism that enables bypassing Spark’s internal data representation and provides the support for vectorized row batches (as proposed in Shen *et al*., 2021) that are used for producing ORC output.

Other auxiliary optimizations including external dependencies configuration and environment setup were evaluated and their impact on the overall performance is described in Results.

### 2.5 Cloud readiness

The increasing availability of cloud computing services for research is gradually changing the way scientific applications are developed, deployed, and run (Vaillancourt *et al*., 2020). To ensure portability and reproducibility of SeQuiLa-based data processing, we followed the Infrastructure as Code (Guerriero *et al*., 2019) and DevOps principles for setting up the computing resources that can be used for both private and public clouds deployments. Hence, we have used technologies like Terraform (for cloud infrastructure provisioning, Modi, 2021), Helm (for deploying applications on Kubernetes clusters, Shah and Dubaria, 2019) and Docker (for application code packaging and shipment, Boettiger, 2015). SeQuiLa has been successfully deployed to both popular managed Hadoop services like Google Dataproc (utilized also in Krissaane *et al*., 2020 and managed Kubernetes services like Google Kubernetes Engine (GKE) or Azure Kubernetes Service. Figure 3 presents an exemplary setup on GKE using the spark-on-k8s-operator and SeQuiLa application defined as a Kubernetes Custom Resource Definition. This architecture was suggested in Castro *et al*., 2019 as preferred Apache Spark deployment scenario for scaling data analytics workloads and enabling efficient, on-demand utilization of resources in the cloud infrastructure. More detailed information on setup and corresponding Terraform modules can be found in the dedicated GitHub repository (https://github.com/biodatageeks/sequila-cloud-recipes).

**Fig. 3.**
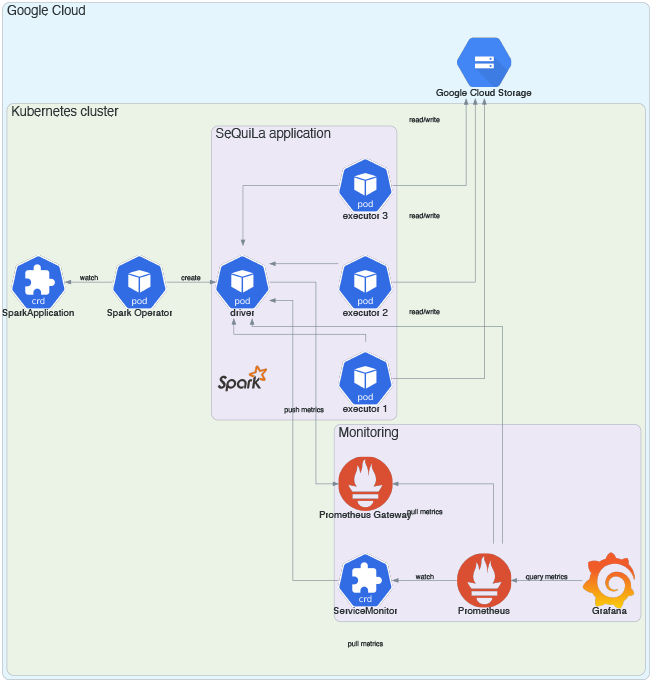
SeQuiLa deployment on GKE with spark-on-k8s-operator with Kubernetes Custom Resource Definition, Prometheus for runtime metrics collection and Grafana as observability platform.

### 2.6 Features

Table 1 summarizes the features of SeQuiLa and compares them with state-of-the-art software including samtools and GATK.

**Table 1.**
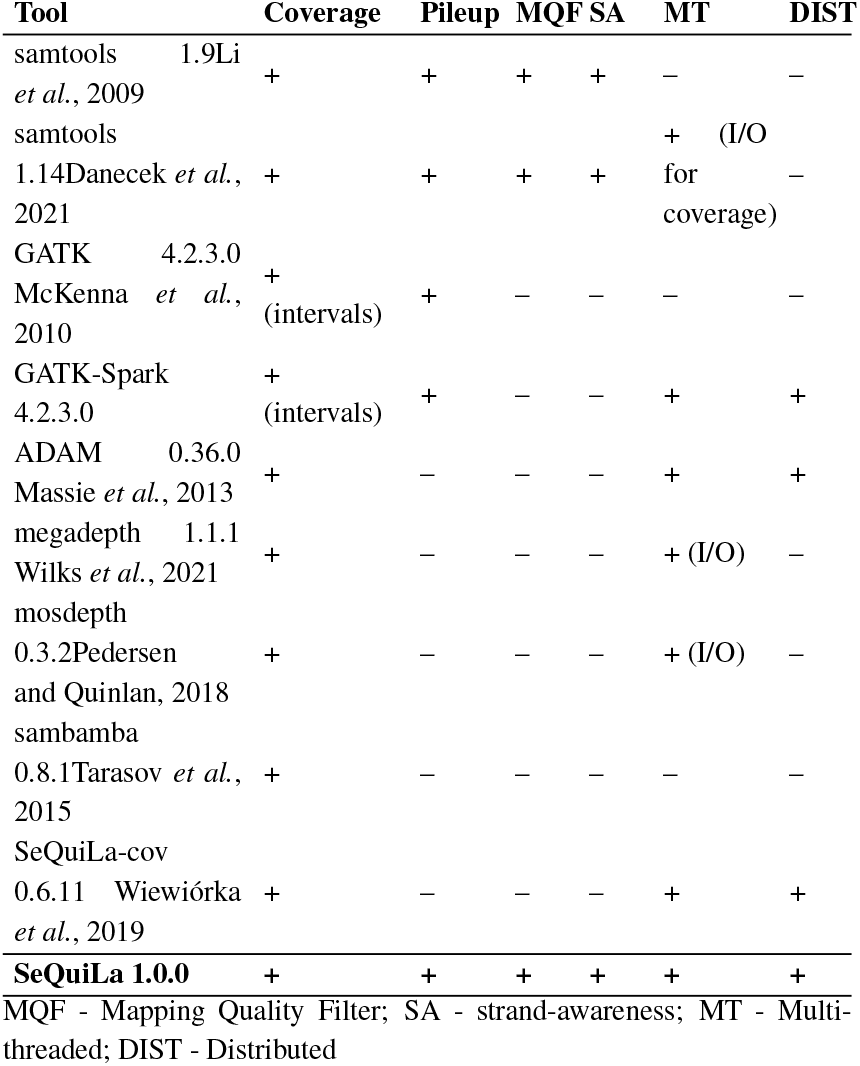
Investigated solutions

SeQuiLa operates on sorted aligned sequencing reads both in BAM and CRAM format. The fast pileup algorithm requires reads to have MD Tag attribute which can be determined during the alignment process or calculated and added to BAM files independently after alignment is completed. MD tag is described in https://samtools.github.io/hts-specs/SAMtags.pdf as a string encoding the mismatched and deleted reference bases, used in conjunction with the CIGAR and SEQ fields to reconstruct the bases of the reference sequence interval to which the alignment has been mapped. This can enable variant calling without requiring access to the entire original reference. Input files can be read either from the local file system, distributed file system, or object storage using a custom data source which allows for representing input reads as relational data. The dataset used for calculating the pileup can be restricted according to the user-provided parameters including reads bit flag and mapping quality.

Samtools and GATK produce verbose output for every coordinate. On the contrary, our software implements lossless block compression of adjacent genomic positions which results in output an order of magnitude smaller. SeQuiLa result includes the genomic coordinates, reference bases, depth of coverage, the ratio of reference to non-reference bases, alternative bases (strand-aware) with occurrences counts, and optionally the base qualities for the positions where at least one non-reference base is present. The output is stored in the popular big data-ready file formats such as ORC or Parquet making it easy to run further analyses e.g. in Apache Spark or tools like Trino (Sethi *et al*., 2019)

SeQuiLa is also distributed as a Python module (pysequila) and can be used on local resources or cloud infrastructure. It is easily integrated with a widespread open-source notebook-based environment for data analysis including Google Colab or Jupyter.

## 3 Results

### 3.1 Datasets

We have used publicly available Exome Sequencing (ES) and Whole Genome Sequencing (WGS) datasets. We performed quality assurance tests on both short reads (sample NA12878) and long reads (guppy), represented in BAM and CRAM formats that were aligned to human reference genome GRCh38 with MD tags included.

### 3.2 Investigated solutions

Table 1 summarizes the functionality of tools included in our comparison. Among the solutions included in the benchmark, only Spark-based GATK and SeQuiLa offer both multi-threaded and distributed versions of the depth coverage and pileup algorithms. For ADAM and SeQuiLa we used Apache Spark 3.1.2 runtime, in the case of GATK that does not provide support for Spark 3.x, we used Apache Spark 2.4.3. Megadepth, mosdepth, and samtools 1.14 are multi-threaded applications but only the parts of their algorithms responsible for the IO operations (BZGF block compression/decompression) are parallelized – the remaining stages of their algorithm are sequential.

We have also included the previous version of SeQuiLa software (0.6.11) to assess the effect of our improved single-pass and cache-less algorithm. Several tools require additional input i.e. genomic intervals in case of GATK’s coverage and PaCBAM’s (Valentini *et al*., 2019) pileup or the list of genomic positions in case of aseq’s (Romanel *et al*., 2015) pileup that restrict processed data and affect algorithm’s computational complexity therefore the aforementioned solutions were not included in the final benchmark.

### 3.3 Testing environment

#### 3.3.1 Single machine

Table 2 presents key information regarding the hardware and operating system configuration of the machine used for benchmark purposes. No hardware or software virtualization has been used.

**Table 2.** Technical specification - single node

| Processor | Base<br>freq<br>[GHz] | CPUs | Total<br>cores<br>(logical) | Memory<br>[GB] | Operating<br>System | Disk |
| --- | --- | --- | --- | --- | --- | --- |
| Intel(R)<br>Xeon(R)<br>E5-2618L v4 | 2.20 | 2 | 20(40) | 256 | RHEL<br>7.8<br>(Maipo) | 3TB<br>(RAID1) |

#### 3.3.2 Hadoop cluster

Hadoop cluster (HDP 3.1.4) consists of 6 master and 34 worker nodes, 680 (1360 logical) cores, 700 TB of Hadoop File System (HDFS) disks, 6.8 TB Yet Another Resource Negotiator (YARN) RAM, and 100 Gbits network. Master node specification was the same as in the case of single-node benchmark, in the case of workers the only difference was in disks configuration - each node has additional 12 disks in Just a Bunch of Disks setup for HDFS storage.

### 3.4 Performance testing scenarios and configuration

We have arranged four testing scenarios: (i) the pileup function performance on the local machine and (ii) its scalability characteristics on the Hadoop Cluster, (iii) the depth of coverage function performance on the local machine, and (iv) its scalability on the Hadoop Cluster. All tools from Table 1 have been included in the presented benchmarks. ES and WGS alignment datasets in the BAM format have been used as inputs. In the case of tools(ADAM, GATK, and SeQuiLa) running on top of Java Virtual Machine(JVM) we used three distributions of Java Development Kit(JDK) – for a single node, we used `GraalVM CE JDK8` for GATK (it does not support JDK11 yet) and `GraalVM CE JDK11` for ADAM and SeQuiLa for running tests on the Hadoop cluster `OpenJDK8` was used for all solutions. For tests using disq library, an additional BAM index has been created.

### 3.5 Results pileup

In pileup benchmarks (Figure 4), SeQuiLa proved to be the fastest tool outperforming samtools in the single thread scenarios by 1.25*x* − 1.4*x* and GATK (both Spark, and non-Spark based) ≈ 3.9 − 6.5*x* GATK (both Spark, and non-Spark based). In the case of Hadoop cluster benchmarks SeQuiLa again proved to be faster by ≈ 2.8 − 5.3*x* than GATK that also required twice as much memory(8 instead of 4GB) per Spark executor to be able to complete the computations. It is worth noting that we were unable to run GATK with 10 or fewer Spark executors(10 cores) facing errors related to too many opened files (even after increasing Linux `nofile` limit to more than 1 million that is more than the recommended value for Hadoop clusters). We have verified that the algorithm’s modularity is gainful as disregarding base qualities further improves SeQuiLa performance by ≈ 35%.

**Fig. 4.**
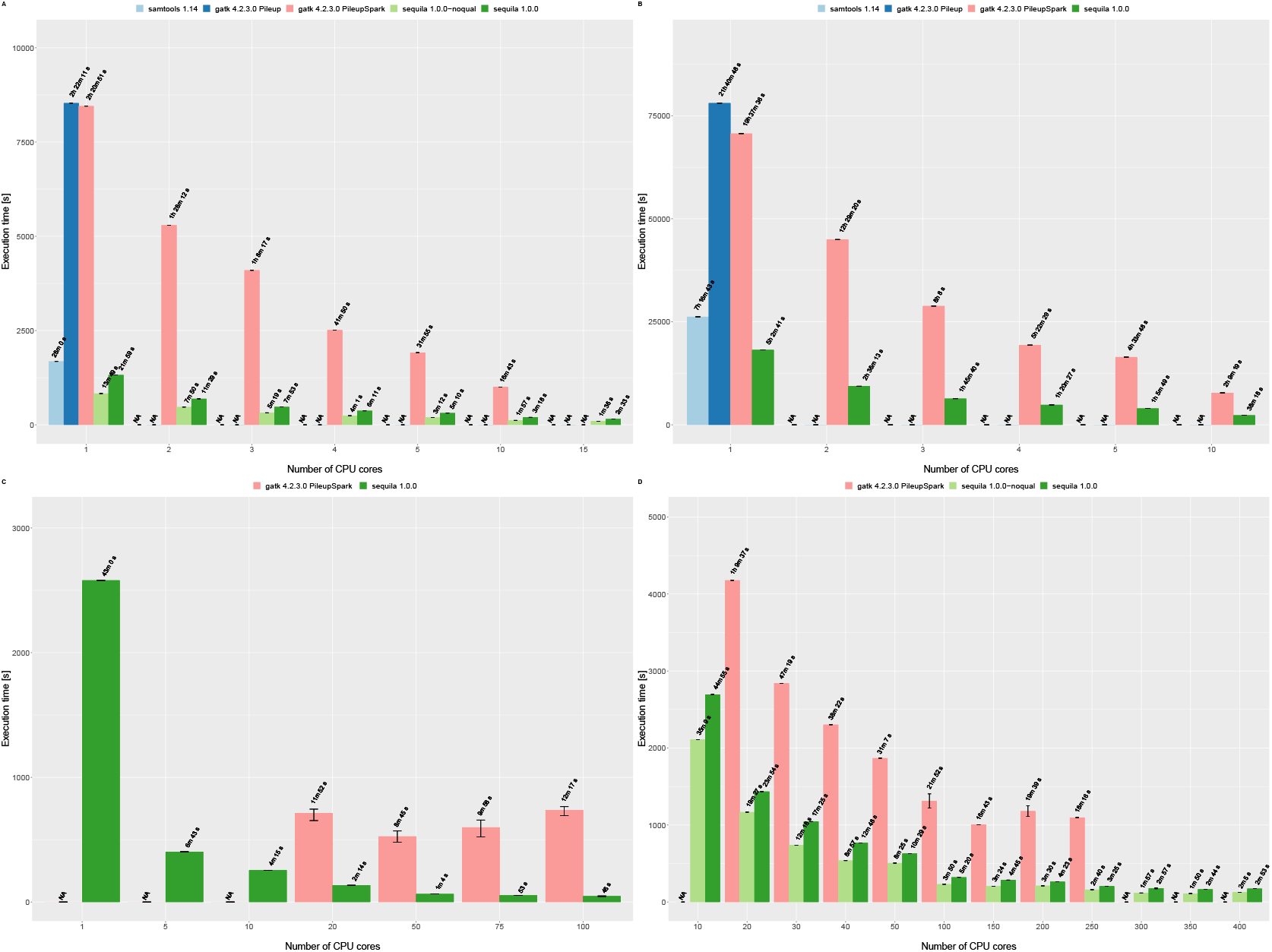
Pileup summary function comparison. Tests were performed on a single node for ES (panel A), WGS (panel B), and on the Hadoop cluster for ES (panel C), WGS (panel D).

#### 3.5.1 Java Virtual Machine optimizations

For development and local testing purposes we have chosen GraalVM which uses an optimized compiler, generating high-performance code, and therefore noticeably accelerates the execution of JVM-based applications (Sipek *et al*., 2020). Additionally, on the source code level, we have applied inlining annotations for frequently called concise methods which are further handled by the Scala compiler, thus avoiding the overhead of method invocation. In our diagnostic tests, we have confirmed that GraalVM choice results in 15% speedup while in-lining improved the timing by another 2% (Figure 5).

**Fig. 5.**
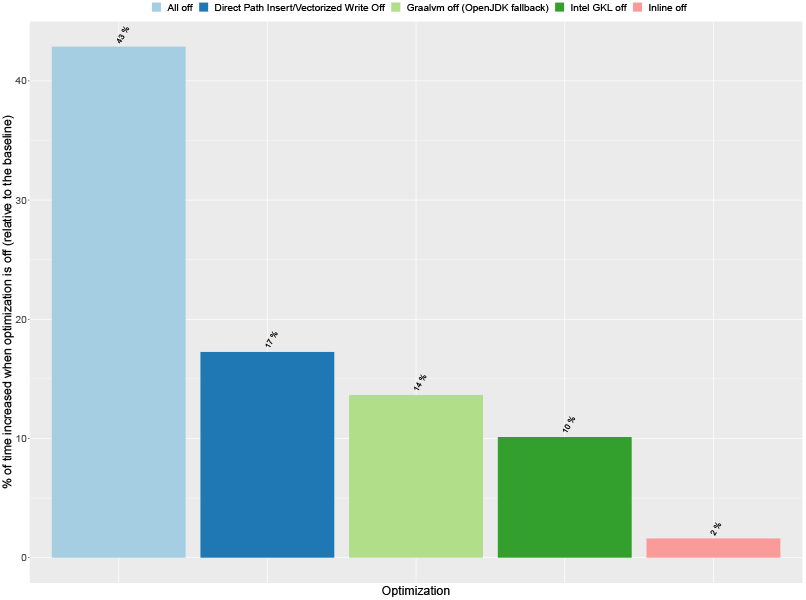
Impact of various optimizations techniques on overall performance as compared with baseline (all optimizations on) for pileup computation

#### 3.5.2 Input-output optimizations

When performing direct vectorized writes into ORC files we have saved 17% of the computing time. We also take advantage of Intel’s Genomics Kernel Library (GKL) providing high-performance operations of decompressing BAM file records. Our benchmarking confirms that the proper use of GKL’s methods results in a 10% decrease of compute time (5).

### 3.6 Results depth of coverage

In the case of both ES and WGS datasets (Figure 6), megadepth proved to be the fastest tool in a single-machine single thread setup outperforming the second one, SeQuiLa by 1.3-1.6x. The gap between them decreases steadily with an increasing number of threads. While processing ES and WGS data SeQuiLa becomes the fastest tool when 5 and 10 threads are used, respectively. It is worth emphasizing that for the following tools: megadepth, mosdepth, and samtools we observed very similar performance characteristics – in contrast to SeQuiLa they do not scale up beyond 5-10 threads at all. These results confirm the fact that these tools only implement the parallel read and blocks decompression operations and the main part of their algorithms does not take advantage of multiple cores. For ADAM we only measured the single-threaded performance that proved to be substantially worse than the remainder of the best performing tools (≈ 40*x*). Last but not least, we confirmed approximately 2x performance improvement in comparison to the first version of the SeQuiLa algorithm initially released 3 years ago.

**Fig. 6.**
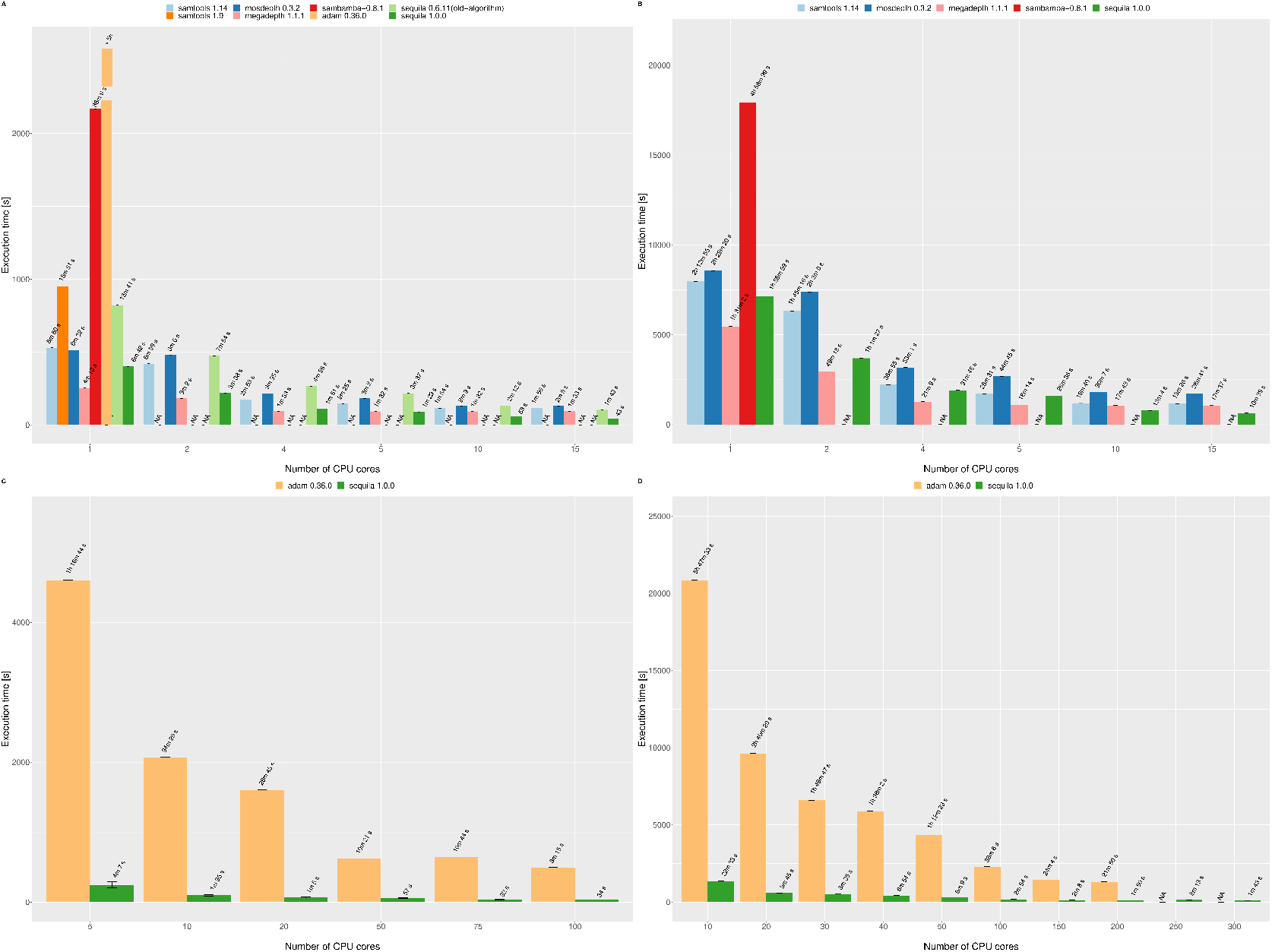
Depth of coverage function comparison. Tests were performed on a single node for ES (panel A), WGS (panel B), and on the Hadoop cluster for ES (panel C), WGS (panel D).

In the case of WGS on the Hadoop cluster (Figure 6) we benchmarked tools allocating from 10 to 200 cores. SeQuiLa outperformed ADAM on average by more than an order of magnitude (11 − 16*x*). We used the same memory (8GB - driver, 4GB executor), Central Processing Unit configurations (1 core) for both Spark processes to ensure comparability of the results between solutions. Also Spark dynamic allocation mechanism has been explicitly disabled. In the case of ADAM tests, we observed random fails of Spark tasks (or even the whole stages) due to network timeouts.

## Discussion

Our pileup method is designed for compatibility with modern distributed computing systems originating from the Hadoop ecosystem mostly implemented using JVM languages such as Java or Scala. This approach incurs additional overheads and causes inefficiencies in the single-node deployments that justify why it does not significantly outperform samtools (a tool written in C language compiling to the native code) in a single thread comparisons. Our pileup algorithm performs especially well on the alignment files with the high-quality short reads when MD tags contain a relatively low number of mismatch/deleted bases. Calculating the complete pileup summaries from the long reads with a large number of mismatches is more challenging for our approach and requires additional modifications that we plan to introduce in its future versions. This limitation does not apply to calculations of the depth of coverage. It also favors BAM over CRAM alignment file formats (data not shown). This is because both the alignment file index scans (random access) as well the sequential reads are much slower (≈ 3 − 4*x*) in the case of CRAM when compared to BAM file format (our results confirms the findings presented in supplementary materials (Bonfield *et al*., 2019). Finally, saving results in the distributed processing can be substantially reduced with adding support for direct, vectorized writes (currently available in the local mode) that is on our project roadmap as well.

Since the complexity of the cloud-native, distributed computing systems have been acknowledged in many studies, including Vaillancourt *et al*., 2020, we have also prepared ready-to-use cloud deployment examples that can help users to start using SeQuiLa in public clouds environments.

## 4 Conclusions

We present a new module that extends and optimizes our SeQuiLa Apache Spark library. This component introducing a new algorithm for fast, scalable, and fully distributed computation of pileup summary from the alignment files (BAM, CRAM). Our solution combines a distributed computing engine based on the extended Apache Spark Catalyst query optimizer with the SQL interface for handling large-scale processing and analyzing next generation sequencing datasets in a consistent tabular form. This approach will help to facilitate the adoption of scalable solutions among users that are neither proficient in distributed computing nor in cloud infrastructures as envisioned in Lawlor and Sleator, 2020.

## 5 Availability of data and materials

The data sets supporting the results of this article are available upon request in Google Cloud Storage bucket: gs://biodatageeks/sequila/data/ (for downloading using Google Cloud Storage compatible tool like gsutil) or https://www.googleapis.com/storage/v1/b/biodatageeks/o?prefix=sequila/data/. Project source code is publicly available (Apache License) at the GitHub platform at https://github.com/biodatageeks/sequila. Cloud deployments documentation and recipes are publicly available at the GitHub platform at https://github.com/biodatageeks/sequila-cloud-recipes. Detailed documentation is available on the project site at https://biodatageeks.github.io/sequila/.

